# Analyzing the neutral and adaptive background of butterfly voltinism reveals structural variation in a core circadian gene

**DOI:** 10.1101/2020.05.13.093310

**Authors:** Olle Lindestad, Sören Nylin, Christopher W. Wheat, Karl Gotthard

**Affiliations:** Department of Zoology, Stockholm University

## Abstract

Many insects exhibit geographic variation in voltinism, the number of generations produced per year. This includes high-latitude species in previously glaciated areas, implying divergent selection on life cycle traits during or shortly after recent colonization. Here, we use a whole-genome approach to genetically characterize a set of populations of the butterfly *Pararge aegeria* that differ in voltinism. We construct a high-quality de novo genome for *P. aegeria*, and assess genome-wide genetic diversity and differentiation between populations. We then use the inferred phylogeographic relationships as the basis for a scan for loci showing signs of divergent selection associated with voltinism differences. The genic outliers detected include population-specific mutations of circadian loci, most notably a locally fixed 97-amino acid deletion in the circadian gene timeless. Variation in timeless has previously been implicated as underlying variation in life cycle regulation in wild populations in our study species, as well as in other insects. These results add to a growing body of research framing circadian gene variation as a mechanism for generating local adaptation of life cycles.

## INTRODUCTION

Organisms in temperate regions often experience a radically different environment depending on the time of year, and adapting life cycles to this seasonality is a major evolutionary challenge. Strong seasonality will favor genotypes that execute particular life cycle events (mating; migration; entering or exiting dormancy) at the appropriate times of year. Because the seasonal cycle is not the same everywhere, but varies with local climate, optimal life cycles also vary geographically. Hence, seasonality provides an opportunity to study divergent adaptation across populations (Conover & Schultz, 1995; Bradshaw et al., 2004; Stinchcombe et al., 2004; Posledovich et al., 2015).

In insects, a central life history trait is voltinism, the number of generations produced by a population in a year. Typically, in a population with one generation per year (univoltinism), every individual at some (species-specific) stage of life enters diapause—a state of hormonally suspended development and metabolic suppression—in anticipation of winter (Tauber et al., 1986; Koštál, 2006). In contrast, a sufficiently long warm season may permit one or several additional, non-diapausing generations to occur each year (bivoltinism or multivoltinism, respectively). Producing a single versus multiple generations per year is a nontrivial difference, as it may be associated with drastic differences in generation time (Roff, 1980), and may necessitate or drive associated adaptations in terms of regulation of diapause induction (Bean et al., 2012; Lindestad et al., 2019), life history traits such as development rate and body size (Masaki, 1972; Fischer & Fiedler, 2002), and developmental pathway expression (Kivelä et al., 2013; Aalberg Haugen & Gotthard, 2015). Nonetheless, many insect species differ geographically in voltinism, typically in relation to the local length of the warm season (Hart et al., 1997; Braune et al., 2008; Hu et al., 2012; Faccoli & Bernardinelli, 2014; Levy et al., 2015).

Expression of a particular local voltinism is often tied to photoperiodism, i.e. how the population responds to differences or changes in daylength. Although quantitative responses in e.g. development rate may play a role (Masaki, 1972; Nylin et al., 1995), the most widely described regulatory mechanism underlying differences in voltinism is local adaptation of the critical daylength below which diapause is induced (Bean et al., 2012; Grevstad & Coop, 2015; Aalberg Haugen & Gotthard, 2015). Critical daylength, across diverse taxa, tends to be locally adapted, often increasing with latitude (Hut et al., 2013). Genetic studies have shown that interpopulation differences in diapause induction are generated by a combination of variation at loci of small and large effects (Ragland et al., 2019). In several cases (Tauber et al., 2007; Yamada & Yamamoto, 2011; Pruisscher et al., 2018; Dalla Benetta et al., 2019), the detected large-effect loci are components of the circadian clock, which exhibit a widely recognized, but controversial and poorly understood, link with photoperiodism across many insect systems (Meuti & Denlinger, 2013). Additionally, allele frequencies at the circadian locus *period* has been shown to co-vary with voltinism in the European corn borer (Levy et al., 2015). Still, the genetic variation associated with interpopulation differences in voltinism in particular remains poorly studied.

The present study explores genetic variation in the speckled wood (*Pararge aegeria*), a woodland-associated satyrine butterfly that varies greatly in voltinism across its pan-European range. The northernmost *P. aegeria* populations are univoltine; southern populations have many overlapping generations per year and appear not to diapause at all (Nylin et al., 1995). Common-garden studies indicate that this life cycle variation is to a large extent generated by local adaptation of photoperiodic plasticity (Nylin et al., 1995; Lindestad et al., 2019). In mainland Scandinavia, *P. aegeria* populations shift from univoltine in central Sweden to bivoltine in southern Sweden and Denmark (Fig. 1a). The southern Swedish populations appear to be recently established—no record of them exists before the 1930s—and are still separated from the central Swedish populations by a gap wherein *P. aegeria* is rare or absent (Nordström, 1955; Eliasson et al., 2005). Bayesian reconstruction from microsatellite data shows that both central Swedish and south Swedish populations likely immigrated from the south via Denmark (Tison et al., 2014), meaning that south Swedish populations went extinct or nearly extinct after the initial northward colonization, and later rebounded either from local stock or through a second migration event via Denmark. In addition, bivoltine populations exist on the Baltic islands of Öland and Gotland, sharing a latitude with univoltine mainland populations. The Öland population is genetically depauperate, while the Gotland population maintains quite high genetic diversity (Tison et al., 2014); the migration histories of both island populations are unknown.

**Figure 1.**
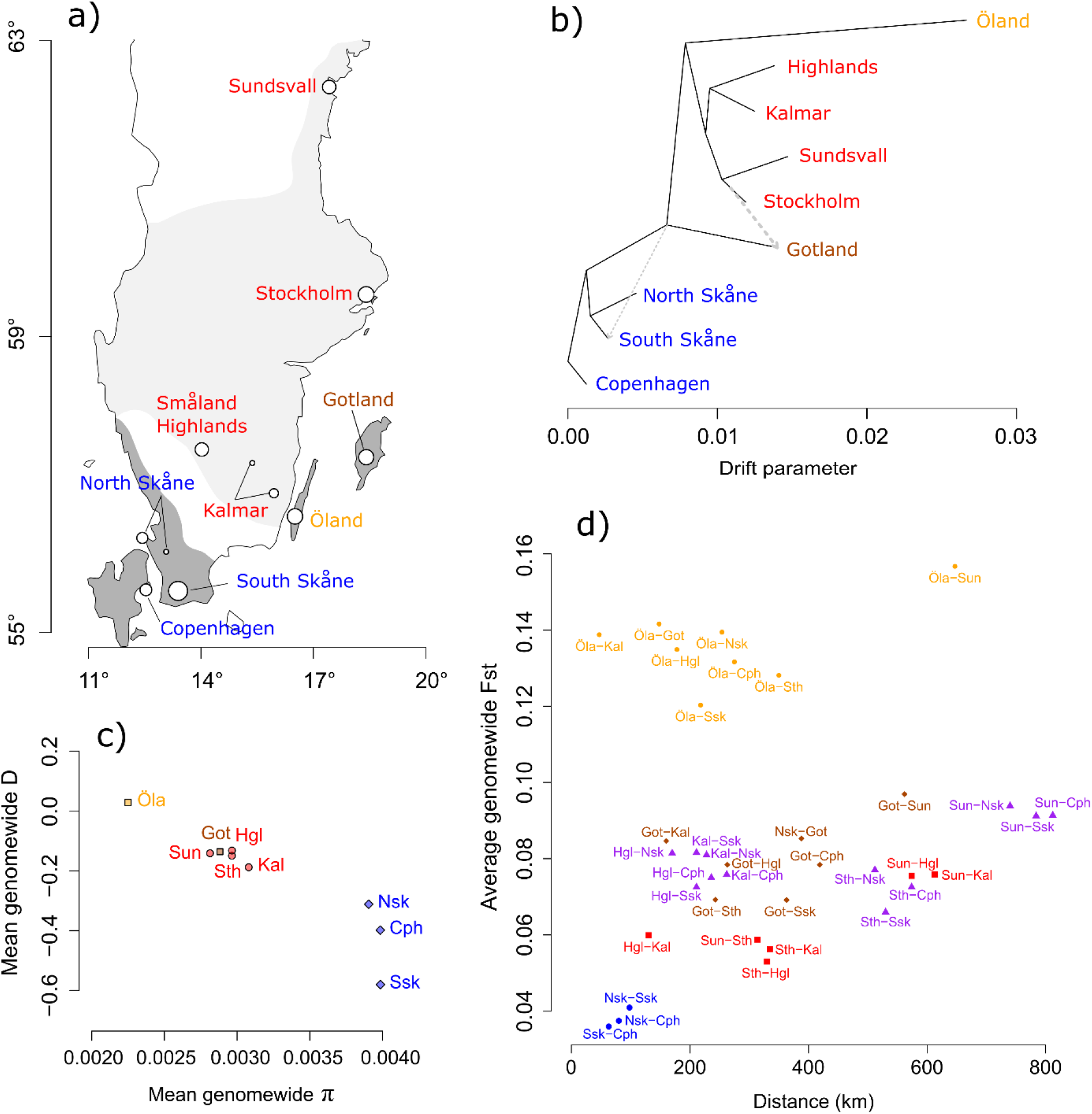
Geographic background and overall patterns of genetic diversity across populations. Except where noted otherwise, northern cluster populations are written in red, southern cluster populations in blue, and the two island populations in orange and brown, respectively. a) Map of *P. aegeria* distribution across Sweden and Denmark. Light-shaded and dark-shaded regions represent predominantly univoltine and bivoltine populations, respectively (adapted from Lindestad et al. 2018). Points mark sampling locations; point size reflects sample size. b) Phylogeographic relationships as inferred by *TreeMix*. Branch lengths represent the amount of genetic drift. Dotted arrows represent inferred migration events; arrow thickness represents the proportion of the recipient population’s ancestry that derives from the donor population. c) Average genome-wide nucleotide diversity (π) and Tajima’s D per population. d) Average genome-wide FST between all pairs of populations. Point shape and color reflects comparison type: large blue circles, southern mainland populations; red squares, northern mainland populations; purple triangles, northern versus southern populations; small orange circles, Öland versus all other populations; brown diamonds, Gotland versus mainland populations.

Previously, a study of genome-wide differentiation patterns between a univoltine (Sundsvall) and a bivoltine population (North Skåne) at either end of the range of *P. aegeria* in Sweden (Pruisscher et al., 2018) identified 15 genomic regions showing signs of divergent selection. SNP variation in these regions—including nonsynonymous substitutions in two circadian loci, *period* and *timeless—*also showed an association with diapause incidence in interpopulation hybrids, indicating that variation at these loci likely contribute to maintaining voltinism variation across Sweden by affecting photoperiodic plasticity. Although SNP genotyping of three additional populations revealed a gradient of allele frequencies at the identified candidate loci across the voltinism cline, much variation remained unexplained. Notably, the Gotland island population is bivoltine and exhibits a relatively low critical daylength (Aalberg Haugen & Gotthard, 2015; Lindestad et al., 2019), despite closely resembling the univoltine populations in terms of variation at the 15 candidate loci (Pruisscher et al., 2018). Here we build on these earlier results in several ways. First, we present the first high-quality genome for *P. aegeria*. Second, we use whole-genome sequence data to reconstruct phylogeographic relationships between nine Scandinavian *P. aegeria* populations, and infer historical gene flow events. Third, we investigate patterns of genome-wide variation among these populations, and scan for genomic regions involved in selection for divergent voltinism. Our findings create a more complete picture of the selective and historical processes underlying variation in voltinism, including a potential contribution from novel variation in circadian genes.

## MATERIALS & METHODS

### Genome assembly and annotation

High molecular weight DNA was extracted from a single *P. aegeria* female from Skåne in southern Sweden, and used for constructing a 10X Genomics Chromium Genome library. Library preparation, sequencing, and genome assembly were performed at the National Genomics Infrastructure, SciLifeLab, Stockholm, Sweden. Genome assembly was conducted using *Super-nova* 1.2 with the phase option enabled; twelve separate assemblies were carried out, each using a different proportion of the 10X sequencing data (from 15% to 100%). Each generated assembly was assessed for two metrics: basic assembly stats, using *Quast* 4.0 (Gurevich et al., 2013), and genic content, using *BUSCO* v. 3 (Waterhouse et al., 2018) with the database *eu-karyota_odb9*. The genome assembly deemed optimal according to these metrics was then scaffolded using a 5-kb insert mate-pair library generated from the same *P. aegeria* individual, after quality filtering and mate pair-specific filtering using *NextClip* v. 1.3. (Leggett et al., 2014), using the scaffolding software *BESST* v. 2.0 (Sahlin et al., 2014). The genome was polished using genomic data from a single individual via the software *Pilon* v. 1 (Walker et al., 2014). Next, the genome was collapsed to a haploid copy using *HaploMerger2* v. 20180603 (Huang et al., 2017), and again assessed for genic content using *BUSCO*, this time with the database *insecta_odb9*. Then, to improve gene model regions, the assembly was super-scaffolded using the *Bicyclus anynana* protein set from Lepbase (Challis et al., 2016) (*Bicyclus_any-nana_nBa.0.1._-_proteins.fa.gz*). Super-scaffolding was carried out using *MESPA* (Neethiraj et al., 2017), which is a wrapper for the high-performing protein-to-genome aligner *SPALN2* (Iwata & Gotoh, 2012). Finally, the genome was annotated using *B. anynana* proteins, as well as a preexisting RNAseq dataset from *P. aegeria*, via the software *BRAKER* (Stankse et al. 2006, 2008, Hoff et al. 2016). Functional annotation was generated using the *eggNOG* v. 5.0 online server with default settings (Huerta-Cepas et al., 2019). Repetitive content was assessed using *Red* (Girgis, 2015).

### Pooled resequencing of populations

DNA for pool-sequencing was extracted from 201 *P. aegeria* individuals sampled from eight populations across Sweden and Denmark (Fig. 1a; Table 1). Most individuals (n=168) were wild adults caught in 2014-2016; of these, 12 female butterflies could not be used, so one offspring each was used instead (the mother was already mated when caught). In order to improve sample size for two of the populations, Öland and Stockholm, 33 individuals were added that had been collected in 2010-2011 for a previous study (Tison et al., 2014). Both sexes were represented for all populations. The tissue used was either half of the thorax (for adults) or the posterior end of the abdomen (for pupae).

**Table 1.**
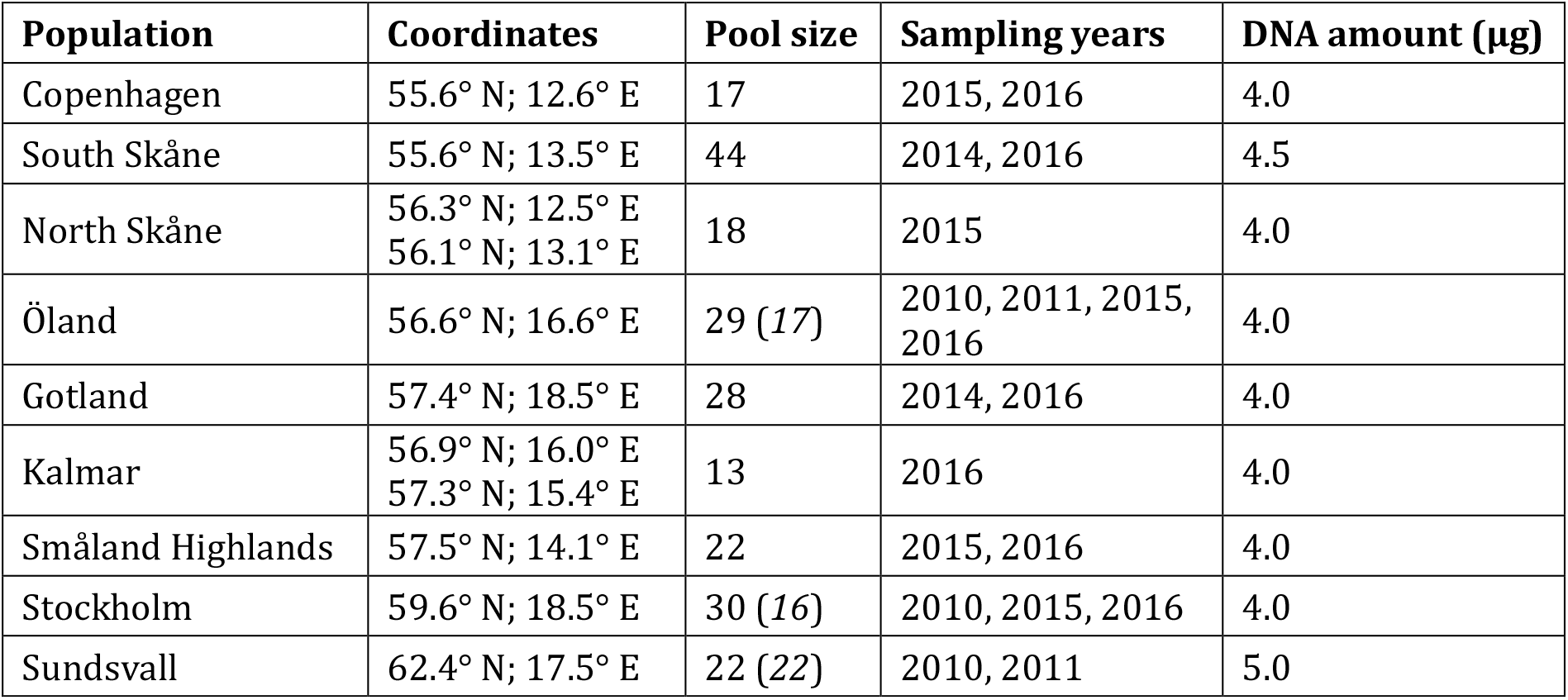
DNA sample details. Pool size is the total number of individuals used; of these, some (numbers in parentheses) were caught for a previous study (Tison et al., 2014). For the Sundsvall pool, DNA extraction and sequencing had also been done previously (Pruisscher et al., 2018). For populations represented by two sampling locations, coordinates for the site contributing the largest part of the sample are listed first.

Extraction was carried out separately per individual, using the standard “cell and tissue DNA” kit on a KingFisher Duo Prime purifier (ThermoFisher Scientific), with added RNAse A to avoid RNA contamination. DNA concentration was measured using a Qubit 2.0 flourometer (ThermoFisher Scientific), and samples were pooled per population using an equal amount of DNA from each individual, yielding 4 – 4.5 μg DNA per population pool. Pooled samples were run on a 2% agarose gel stained with GelRed to visually ascertain that DNA fragmentation was minimal. Library preparation and sequencing (Illumina HiSeq) was performed by SciLifeLab (Uppsala, Sweden), using 150-bp paired-end reads with 350 bp insert size.

In addition to the eight population read sets generated here, an additional *P. aegeria* read set (100 bp paired-end reads; 450 bp insert size) from Sundsvall in northern Sweden was used in our analyses. The Sundsvall dataset was generated from 22 adult *P. aegeria* individuals caught in 2011 for a previous study, using similar extraction, pooling and sequencing methods (Pruisscher et al., 2018).

### Dataset preparation

Raw sequencing files were filtered for PCR duplicates using the *clone_filter* script of *Stacks* 1.21 (Catchen et al., 2013), and Illumina sequencing adapters were removed using *BBDUK2* (Bushnell, 2015). All nine PoolSeq read sets were mapped to the *P. aegeria* genome using *NextGenMap* v. 0.4.10 (Sedlazeck et al., 2013) at 90% identity. Alignments were filtered using *SAMtools* v. 1.6 (Li et al., 2009), keeping only properly paired reads with a map quality of 20 or higher. Read depth (RD) exceeded pool size over large parts of the alignments, leading to a risk of unequal read sampling of sequenced individuals, and hence skewed estimates of nucleotide diversity (Hoban et al., 2016). Because parts of the analyses relied on accurate diversity estimates, and for consistency across analyses, we applied a subsampling approach. First, each alignment was converted into a population-wise pileup file using *SAMtools*. To avoid spurious SNP calling, indels were filtered out (with a 5-bp margin on each side) using scripts from the *PoPoolation* v. 1.2.2 package (Kofler, Orozco-terWengel, et al., 2011). Next, also using *PoPoolation*, reads were subsampled without replacement from each pileup up to a given target RD; pileup positions where raw RD was lower than target RD were excluded from analysis. The 5^th^ percentile of raw RD for each population (or a minimum of 20) was used as the target RD. Likewise, pileup positions in the 99^th^ percentile of per-population RD were excluded. These nine subsampled population-wise pileups (details in Table S1) served as the input for all down-stream applications. A phred quality score cutoff of 20, and a minimum count of 2 for SNP alleles, were used in all analyses.

### Population history and connectivity analyses

To estimate overall population differentiation, the average genome-wide fixation index (F_ST_) was calculated for all possible population pairs using *PoPoolation2* v. 1.201 (Kofler, Pandey, et al., 2011). All nine input pileups were joined together as separate columns in a single mpileup file, which was then converted into the sync format. SNPs were called on this sync file, yielding about 7.8 million SNPs. F_ST_ was calculated on the called SNP variation in nonoverlapping 5000-bp windows across the whole genome. The mean F_ST_ across all windows for each pairwise population comparison was then calculated. Here and elsewhere in the analyses, F_ST_ was calculated from allele frequencies as per Hartl & Clark (2007). Additionally, overall genetic diversity in each population was surveyed by measuring nucleotide diversity (π) and Tajima’s D across each single-population pileup, using *PoPoolation*. The D statistic measures the relationship between nucleotide diversity and the number of variable sites, and can provide information on demographic history (Tajima, 1989). For both the F_ST_, π and Tajima’s D analyses, only genomic windows where at least 50% of positions met the RD requirements were analyzed.

In order to investigate population history and potential gene flow, a phylogeographic analysis was conducted using *TreeMix* v1.13 (Pickrell & Pritchard, 2012). *TreeMix* takes genome-wide allele frequency distributions as input, reconstructs a bifurcating phylogeny, and identifies population pairs which share more allele frequency variation than the reconstructed tree can account for. The software then allows these incongruences to be resolved by sequentially adding inferred migration events between points on the tree, where each migration event is modelled as a unidirectional donation of a specific percentage of a population’s allelic variation. Input allele frequencies were extracted from the same nine-population sync file as for the F_ST_ analysis. *TreeMix* was run five times, each time allowing for one additional inferred migration event, from zero migration events to four. Because previous results indicate that all Swedish regions were settled from the south (Tison et al., 2014), Copenhagen was set as the root population in all runs.

### Scanning for outlier regions

Based on the results of the phylogeographic analyses (Fig. 1), we set up three different comparisons between a bivoltine and a univoltine region. Here, the univoltine Sundsvall, Stockholm, Kalmar and Småland Highlands populations were treated as a single phylogeographic unit (hereafter “northern cluster”). In order to analyze this cluster as a single unit, we merged the input pileups using a custom *awk*-based *bash* script that concatenated the base calls from the constituent populations at each genomic position, generating a single base call column representing all reads for this cluster. Likewise, Copenhagen and the two Skåne populations were treated as a single bivoltine population (hereafter “southern cluster”), and their pileups combined in the same way. The three paired comparisons, then, were Öland vs the northern cluster, Gotland vs the northern cluster, and the southern cluster vs the northern cluster. The aim was to find genomic regions showing (1) high differentiation between two populations of differing voltinism, and (2) relatively low genetic diversity, which together may indicate the past occurrence of a divergent selective sweep (Maynard Smith & Haigh, 1974; Carneiro et al., 2014; Reid et al., 2016).

For each comparison, patterns of genetic variation (π and F_ST_) were calculated across the genome, again in nonoverlapping 5-kb windows. π was calculated using *PoPoolation* v1.2.2 on individual pileups for each geographic region in the pair; F_ST_ was calculated using *PoPoolation2* v1.201 on a single mpileup of all four regions (where the northern and southern clusters had been merged into a single base call column each, as described above). Finally, in addition to the sliding-window analyses, F_ST_ was calculated in pairwise mpileups on a nucleotide-by-nucleotide basis, i.e. for each individual SNP in the genome. Biallelic SNPs with an F_ST_ of 0.9 or higher were considered strongly differentiated (when two populations are completely fixed for different alleles of a SNP, F_ST_ = 1).

Outlier windows were selected based on four criteria: (1) an overall window F_ST_ in the 99^th^ percentile of windows for that comparison, (2) nucleotide diversity in the 1^st^ percentile of windows for either population in that comparison, (3) containing at least one strongly differentiated SNP, and (4) being located in a genic region. The latter criterion was assessed using *BEDtools* v2.21.0 (Quinlan & Hall, 2010), by cross-referencing the list of potential outlier regions with the genic regions identified in the genome annotation. A genic region was defined as all introns and exons of a putative gene, plus the 5 kb on either side (i.e. immediately upstream and downstream), to account for potential selection on regulatory mutations. Each hit region was inspected visually for placement of differentiated SNPs, etc. using the Integrative Genomics Viewer (Robinson et al., 2011), and the identity of the gene(s) overlapping with each outlier region was checked by running the putative amino acid sequences against the NCBI database using BLASTp (Camacho et al., 2009), specifying Lepidoptera as the focal taxon.

### Scanning for population-specific indels

Because one of the identified outlier genes contained a large, population-specific deletion, a follow-up analysis was conducted in order to assess how common this type of structural variation is in the *P. aegeria* genome. For this analysis, the read depth-filtered dataset was not used, so as not to unnecessarily lose data; instead, a single mpileup file was generated for all nine populations. Positions where read depth equaled zero for all populations were filtered out, and an indel scan was then run in 100-bp windows using a custom *awk*-based *bash* script. Each 100-bp window where a single population had zero reads, while all other populations showed an average read depth of 20 or more across the window, was scored as an indel. Note that using this method, potential indels varying within a population, or shared between more than one population, were not counted.

## RESULTS

### *P. aegeria* genome assembly

A total of 148.12 Mreads (85% ≥ Q30) were generated during sequencing, and using different proportions of this raw read data for assembly greatly affected assembly statistics (Fig. S1). For the 12 initial assemblies generated, N50 values ranged from 1.8 to 16.9 kbp, while total assembly length ranged from 198.1 to 466.7 Mbp. The 100% and 35% assemblies showed the highest N50 values (16.9 and 14.2 kbp, respectively), as well as the largest number of scaffolds at least 50 kbp in length (n=778 and 661, respectively). The 35% assembly also showed superior gene content and quality, with 74% of BUSCO entries being full length and single copy, and the lowest number of missing orthologs (7.3%). Given these results, the 35% assembly was used as the backbone for subsequent scaffolding, polishing, haplomerging, protein scaf-folding and annotation. The final genome had an N50 of 513 Mbp, consisting of 26,567 scaffolds, with 95 scaffolds > 1 Mbp, a maximum scaffold length of 3,041,493 bp, and a total of 10.9% non-ATGC characters. BUSCO analysis of the final genome was much improved, with 88% entries complete and single-copy, 5.7% fragmented, and 5.7% missing. A total of 23,567 high-quality genes were annotated, of which 15,993 with functional annotations, and 33% of the genome was found to contain repetitive content.

### Population history and connectivity analyses

Phylogeographic analysis suggested a deep split between southern and northern/island *P. aegeria* populations, with the two Skåne populations as the sister group to all other Swedish populations, and all four northern univoltine populations clustering together (Fig. 1b). At least two admixture events could be inferred from the residual allelic variation. Firstly, Gotland appears to derive a large part of its genetic variation (~21%) from a population related to the Stockholm population. Secondly, Southern Skåne is inferred to have received a smaller amount of genetic variation (~5%) that is shared by all northern and island populations. Subsequent migration edges were not considered geographically likely to reflect true gene flow events, and did not notably improve tree fit, so were not considered (Fig. S2).

Average genome-wide F_ST_ varied between population comparisons (Fig. 1d). Mainland populations showed a weak isolation-by-distance pattern, with low differentiation between the three closely grouped southern (Copenhagen and Skåne) populations, and generally higher differentiation between the northernmost population (Sundsvall) and other populations. However, most comparisons across the north/south distribution gap gave an F_ST_ around 0.07-0.08, regardless of geographic distance. As for the two island populations, Öland was relatively strongly differentiated (F_ST_ ≈ 0.14) from all other populations, while Gotland was mildly differentiated (F_ST_ ≈ 0.08) from all mainland populations. In terms of population-wise patterns of genetic diversity, the three southern populations were relatively diverse, with a tendency toward negative values of Tajima’s D, while the northern mainland populations showed intermediate diversity and less negative values of D. Again, the two island populations yielded quite different results: Öland had low nucleotide diversity and near-zero average Tajima’s D, while Gotland showed similar patterns of diversity to the northern mainland populations (Fig. 1c).

### Genic outlier regions

All three univoltine/bivoltine comparisons generated similar numbers (n=22–28) of genic regions showing high F_ST_ and low π, but the results differed strongly when the search was limited to windows containing at least one strongly differentiated SNP (Fig. 2). All outlier SNPs detected were unique to a single comparison (Table 2), and the discovery rate corresponded to the overall differentiation between the two geographic regions in the pair. When comparing the southern cluster to the northern cluster (Fig. 2e-f), overall genome-wide F_ST_ was low (mean=0.04), and only one region met all of the search criteria: a putative glucose dehydrogenase gene, which contained a nonsynonymous exonic substitution. Several of the outlier loci detected in Pruisscher *et al’*s (2018) scan of two of the populations represented in this comparison (Sundsvall and North Skåne) also showed a high F_ST_/low π signal here, including the circadian genes *timeless* and *period*. However, the previously described candidate SNPs in these genes did not register as strongly differentiated here, as the northern Kalmar population harbored the “southern” alleles at intermediate frequencies. By far the largest number of outlier SNPs was found when comparing Öland to the northern cluster: 13 windows met all search criteria (Fig. 2c-d), although two of the putative gene sequences, which were very short and gave no BLASTp result, may represent annotation artefacts. Three outlier loci contained non-synonymous substitutions: *kinesin-associated protein 3*; a calcium-binding protein; and a post-transcriptional methylase subunit. Meanwhile, the Gotland/northern cluster comparison yielded outlier windows in three genes. The first was a programmed cell death protein; the latter two were *timeless* and *period*, both of which contained SNPs unique to Gotland, including a nonsynonymous (Pro→Ser) substitution in *period*.

**Table 2.**
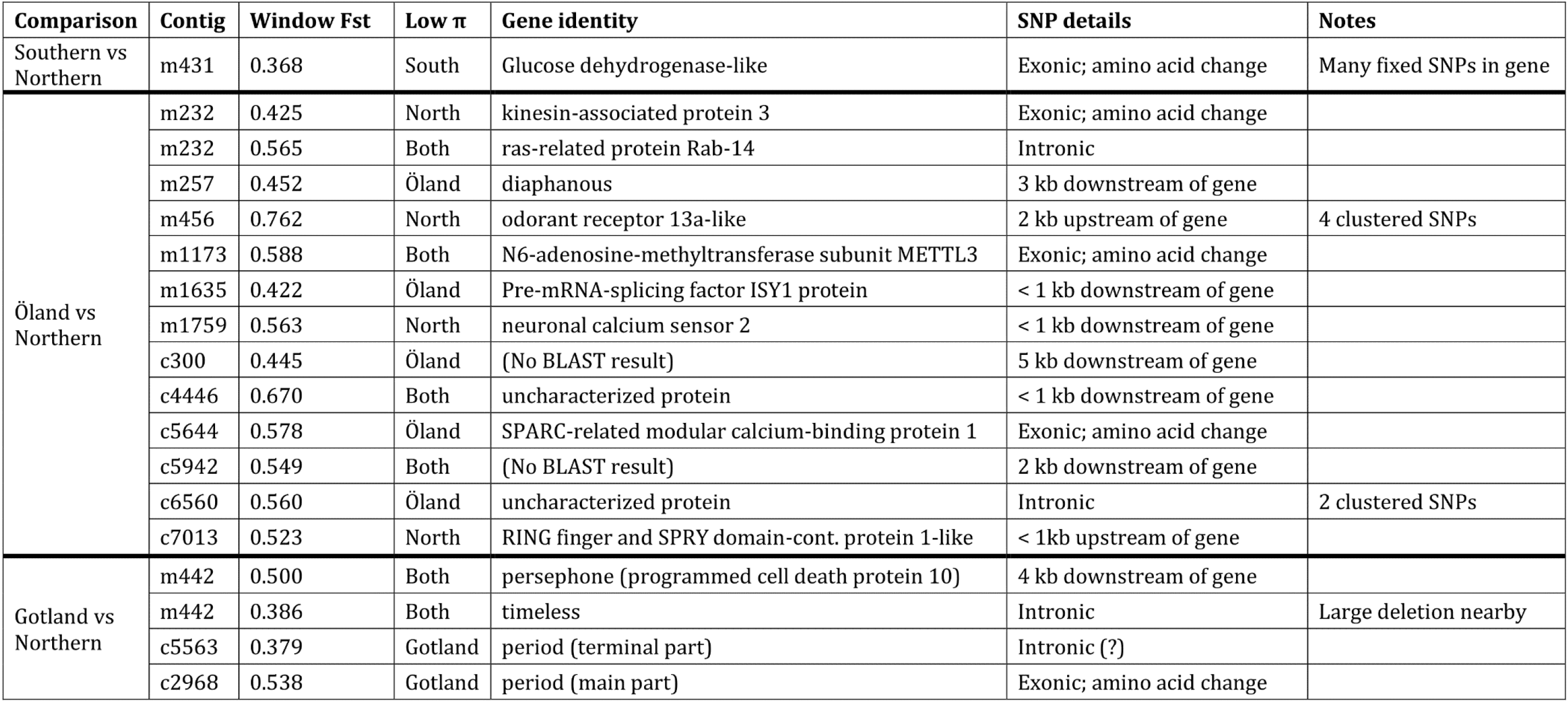
Details on genic outlier SNPs detected for each comparison between a bivoltine region/population and the univoltine northern region: F_ST_ for the focal 5000-kb outlier window, which population(s) showed a low local value of π, gene identity according to NCBI BLAST, and details on the location of the strongly differentiated SNP(s). Note that *period* appears to have been split across multiple contigs in the genome assembly; two of these contigs registered as outliers. As the SNP found in contig 5563 lay upstream of the gene fragment matching the terminal end of *period*, it is assumed to in fact be intronic.

**Figure 2.**
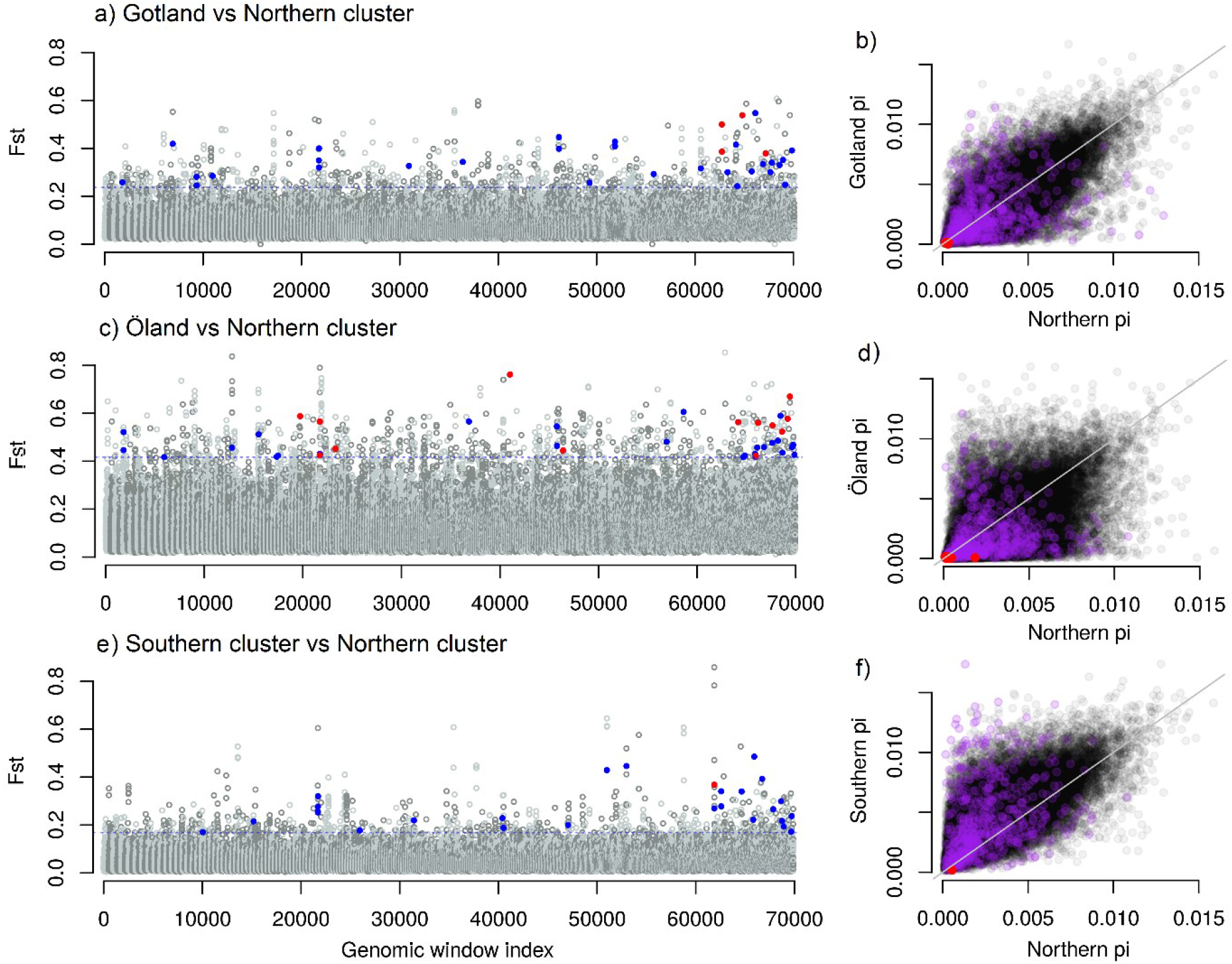
Results of genetic outlier scan. Left panels (a; c; e): Manhattan plots of differentiation between pairs of geographic regions with differing voltinism. Each point represents a genomic window 5000 bp wide; contigs are sorted by length and colored alternately light and dark grey. Dotted line: 99^th^ percentile of F_ST_ for this comparison. Genic outliers with high F_ST_ and low nucleotide diversity are highlighted in blue; outliers also containing strongly differentiated SNPs are highlighted in red. Right panels (b; d; f): relationship between genome-wide nucleotide diversity for either population in a pair, with unity line. 5-kb genomic windows showing an F_ST_ above the 99^th^ percentile are colored purple; remaining windows are colored black. Genic SNP outliers are again marked in red.

Additionally, visual inspection of the alignments revealed an otherwise high-coverage region of approximately 1500 bp in *timeless* where no Gotland reads had mapped, indicating a deletion unique to this population (Fig. 3). This deletion spanned exon 15 (out of 16) of *time-less* as well as most of exon 14, shortening the predicted TIM protein sequence by 97 amino acids, or 8% of its length. A genome-wide scan of population-specific indel variation revealed putative deletions at an additional 325 locations in the genome, some of a similar size, but most only one or two hundred bp in size (Fig. S3). In 96 cases, the missing sequence data lay within an exon. The majority of putative deletions were in Sundsvall, although that population may have a somewhat higher risk of false positives, as its read depth was generally lower across the genome. Öland showed deletions at a few loci, and Gotland at one additional locus. However, the deletion in *timeless* was the largest population-specific indel detected.

**Figure 3.**
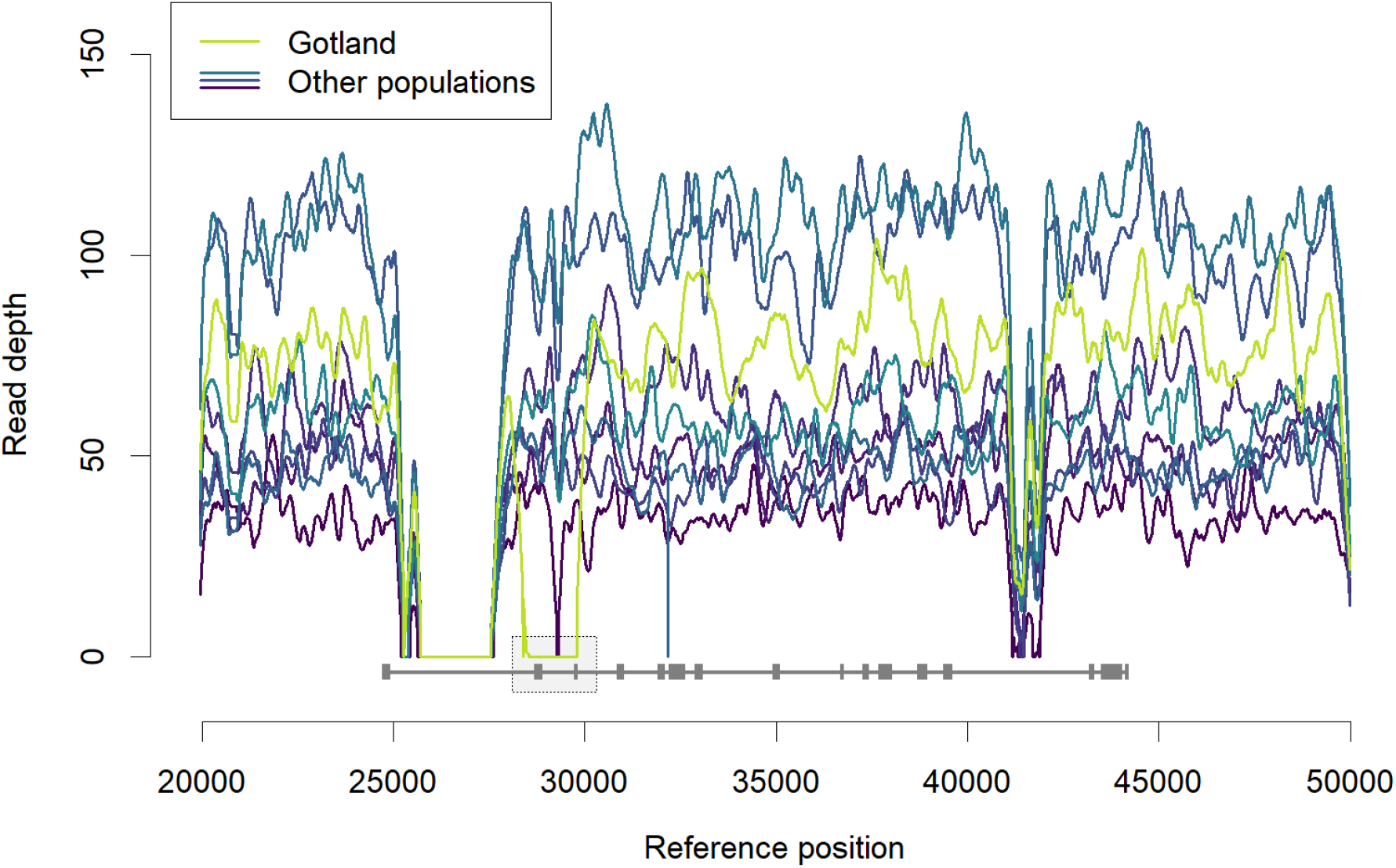
Read depth across *timeless* for Gotland (bright green) versus all other populations (blue). All non-zero values have been smoothed using a 400-bp moving average. The grey line shows the inferred *timeless* transcript, with introns as narrow sections and exons as thick sections (the first exon is rightmost; the sixteenth and terminal exon is leftmost). The dotted rectangle highlights a region from approximately 27,000 to 28,000 bp where Gotland read depth falls to zero, indicating a population-specific deletion that spans exon 15 and most of exon 14. Other low-RD regions are shared between populations, and probably represent repetitive elements.

## DISCUSSION

Scandinavian *P. aegeria* constitute a set of relatively recently (i.e. postglacially) immigrated populations showing a certain degree of local adaptation to a climate gradient. Consistent with this, the results presented here suggest a genetic landscape of mild interpopulation differentiation, interspersed with spots of stronger divergence with potential adaptive importance. The outlier search was conducted with unspecific criteria, the aim being to identify any genic regions potentially associated with selection on voltinism variation. Given this, it is notable that additional nonsynonymous variation was uncovered in both *timeless* and *period*, two representatives of the core set of circadian loci previously implicated in generating variation in life cycle regulation in *P. aegeria* (Pruisscher et al., 2018) as well as other species (Tauber et al., 2007; Yamada & Yamamoto, 2011; Paolucci et al., 2016). The exonic deletion in *timeless* was found essentially by luck, as our methods included a visual examination of each detected outlier region, revealing the local drop in read depth (Fig. 3). Judging by follow-up scans, such large, population-specific deletions are not particularly common in *P. aegeria* (Fig. S3). The chance nature of its discovery highlights the risk, inherent to typical SNP-based genomic methods, of overlooking structural variation that may be important to adaptation (Hoban et al., 2016).

Overall genetic differentiation among the studied populations was fairly low, with most comparisons yielding an average genome-wide F_ST_ of 0.06 – 0.08 (Fig. 1d). The exception was Öland, which showed relatively strong differentiation from all other populations; this likely results from the low allelic diversity of this population (Fig. 1c), which appears to have gone through a demographic bottleneck during its history, as also indicated by its long branch in Fig. 1b. On the mainland, both *TreeMix* and genome-wide F_ST_ results indicate two well-separated population clusters with differing voltinism. Gene flow across the mainland voltinism divide is probably limited or absent, given that the two southernmost univoltine populations,

Kalmar and the Småland Highlands, are just as differentiated from the bivoltine populations as Stockholm is, at twice the geographical distance (Fig. 1d). In general, regions of elevated F_ST_ between the compared populations showed no particular tendency toward lowered nucleotide diversity in either population (Fig. 2b; d; f). This differs strongly from results obtained with similar methods from a study system where populations diverged adaptively on a much more recent timescale (Reid et al., 2016), and indicates that the largest part of genome-wide differentiation between Scandinavian *P. aegeria* populations has occurred through neutral genetic processes over the course of postglacial migration, rather than through selection. Overall nucleotide diversity was higher for southern populations, which confirms earlier microsatellite results (Tison et al., 2014) and is consistent with gentle clinal loss of diversity under northward migration from glacial refugia. Interestingly, low average π values correlated with high values of D (Fig. 1c). This may reflect a relatively short time having passed since the migrational founder event, with rare variants having been lost to drift and not yet replaced by novel mutations.

The *TreeMix* analysis suggests that South Skåne (but not North Skåne) has received a small amount of novel genetic variation shared by all northern and island populations. This could be interpreted as gene flow along the Baltic coast. However, a more likely explanation may be secondary contact, where the recent (i.e. 1930s) migration wave from Denmark encountered a small number of *P. aegeria* still present in southeastern Sweden from the first migration wave that produced the northern/island populations. Our analyses also suggest a possible explanation for the surprisingly high genetic diversity of the geographically isolated island of Gotland (Fig. 1c), namely that this population may have originated through admixture of different founder populations. The results of these phylogeographic analyses are fully compatible with, and contribute additional detail to, the picture of genetic diversity in Northern European *P. aegeria* suggested by earlier results (Tison et al., 2014).

Given the differentiation in both voltinism regulation (Nylin et al., 1995; Lindestad et al., 2019) and associated life history traits (Aalberg Haugen & Gotthard, 2015) displayed by Scandinavian *P. aegeria* populations, one of the aims of the present study was to investigate the extent to which selective (local adaptation) versus non-selective processes (genetic history) have contributed to the life cycle patterns expressed. Selective and historical processes may both shape population traits (Hereford, 2009), depending in part on how closely tied the measured trait is to fitness (Travisano et al., 1995). A particularly good indicator of selection is functional differentiation between populations despite a certain degree of gene flow, as it suggests that phenotypic differentiation is continually being maintained by local adaptation (Kawecki & Ebert, 2004). Given that univoltine and bivoltine populations clustered separately in the phylogeographic analyses, with little evidence of recent or ongoing genetic exchange between northern and southern mainland populations, such conclusions cannot easily be drawn here. However, it should be noted that voltinism is a labile trait on ecological timescales (Altermatt, 2010), generated through a combination of genetic and environmental differences between populations (Lindestad et al., 2019), and has been observed to evolve quickly through modification of photoperiodic plasticity (Yamanaka et al., 2008; Bean et al., 2012). For these reasons, a classical phylogenetic-comparative interpretation with a single transition of voltinism states along the population tree may be misleading. Assuming a postglacial colonization scenario where a univoltine range margin (as today represented by Sundsvall) tracks the upper limit of the viable climate envelope for the species, it is likely that currently bivoltine *P. aegeria* populations in this region were univoltine when first established. Interestingly, the bivoltine Gotland population, which in terms of the candidate loci for diapause regulation characterized by Pruisscher *et al* (2018) has a distinctly “northern” profile, is here shown to harbor additional, unique variants at two of these same candidate loci. This could be interpreted as a result of selection for bivoltinism through a lowered diapause induction threshold following colonization by univoltine genetic stock. It is also worth noting that the Gotland population appears to have an intrinsically longer diapause than neighboring populations, possibly as an adaptation to partial bivoltinism (Lindestad et al., 2020). Differences in the rate of diapause termination are associated with variation at *period* in the moth *Ostrinia nubilalis* (Levy et al., 2015; Wadsworth & Dopman, 2015), marking the novel circadian gene variation in Gotland as a possible parallel.

Local alignment of the predicted *P. aegeria* TIM protein sequence with that of *Drosophila melanogaster* shows that the Gotland-specific deletion overlaps at least partially with the cytoplasmic localization domain (CLD) characterized by Saez & Young (1996). In *D. melanogaster*, this domain prevents TIM from entering the nucleus; its effect is overridden by dimerization of TIM with PER, which allows the transcriptional feedback loop of the circadian clock to be closed. Inferring functional consequences of circadian gene variation across insect orders is problematic, not least because the circadian clock machinery is variable. In Lepidoptera (moths and butterflies), PER and TIM appear not to play the exact same roles as in drosophilids (Sauman & Reppert, 1996; Zhu et al., 2005; Yuan et al., 2007), although the protein sequences of both TIM and PER in domestic silkmoth do retain their respective interaction sites (Iwai et al., 2006). Nonetheless, the overlap of the deletion with these putative domains is an indication that the deletion in Gotland *P. aegeria* may have some effect on the functioning of the TIM protein.

European populations of *P. aegeria* are a useful model system for studying postglacial migration and climate adaptation (Hill et al., 1999; Tison et al., 2014; Aalberg Haugen & Gotthard, 2015; Pateman et al., 2016). The present results expand on earlier analyses in this species by providing a more fine-scale picture of both neutral and putatively adaptive variation. In a wider perspective, it is especially notable that circadian loci, which appear to play a key role in adaptation to geographic variation in seasonality across a range of insect species, again emerge as potentially under selection. While any hypothetical effects of the novel mutations described here (either on circadian functioning or photoperiodic diapause induction) are unknown, these loci stand out as promising targets for further investigation *in vivo*.

## ACKNOWLEDGEMENTS

This work was financed by the Knut and Alice Wallenberg Foundation (KAW 2012.0058), the Bolin Centre for Climate Research at Stockholm University (to K. Gotthard), and the Swedish Research Council grants VR-2012-3715 (to S. Nylin) and VR-2017-04500 (to K. Gotthard). Many thanks to Maria de la Paz Celorio Mancera for help with DNA extraction and sample preparation, and to Peter Pruisscher for input on bioinformatic methods.

## Notes

### Competing Interest Statement

The authors have declared no competing interest.

