## Supplementary tables and figures for "Analyzing the neutral and adaptive background of butterfly voltinism reveals structural variation in a core circadian gene"

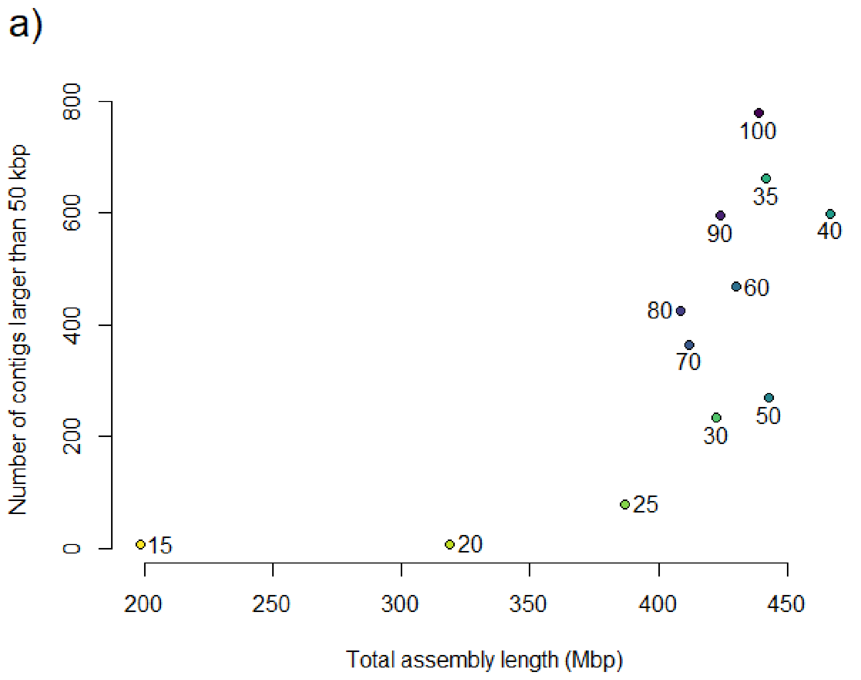


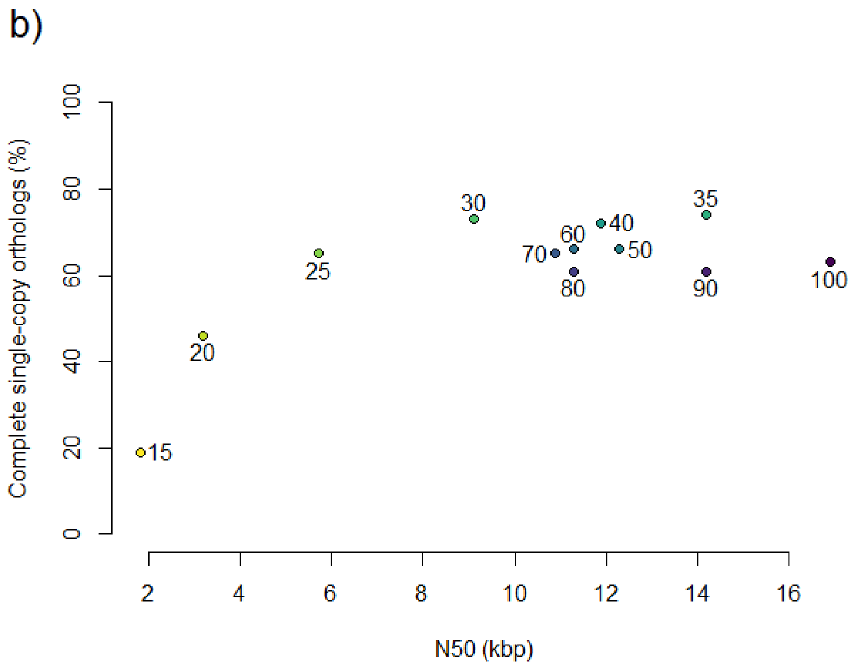


**Figure S1.** Assembly performance (size, fragmentation and genic content) of the 12 initial *P. aegeria* genomes generated. Point labels refer to the percentage of raw read data used to generate each assembly. **a)** Total assembly length versus number of contigs larger than 50,000 bp. **b)** N50 score (a measure of assembly fragmentation) versus the proportion of the 303 single-copy orthologs in *BUSCO*'s *eukaryota_odb9* catalogue that were found in complete form. The 35% assembly was used as the base for further genome construction, as it combined low fragmentation, and a total size similar to known Lepidopteran genome sizes, with a large number of high-quality gene models.

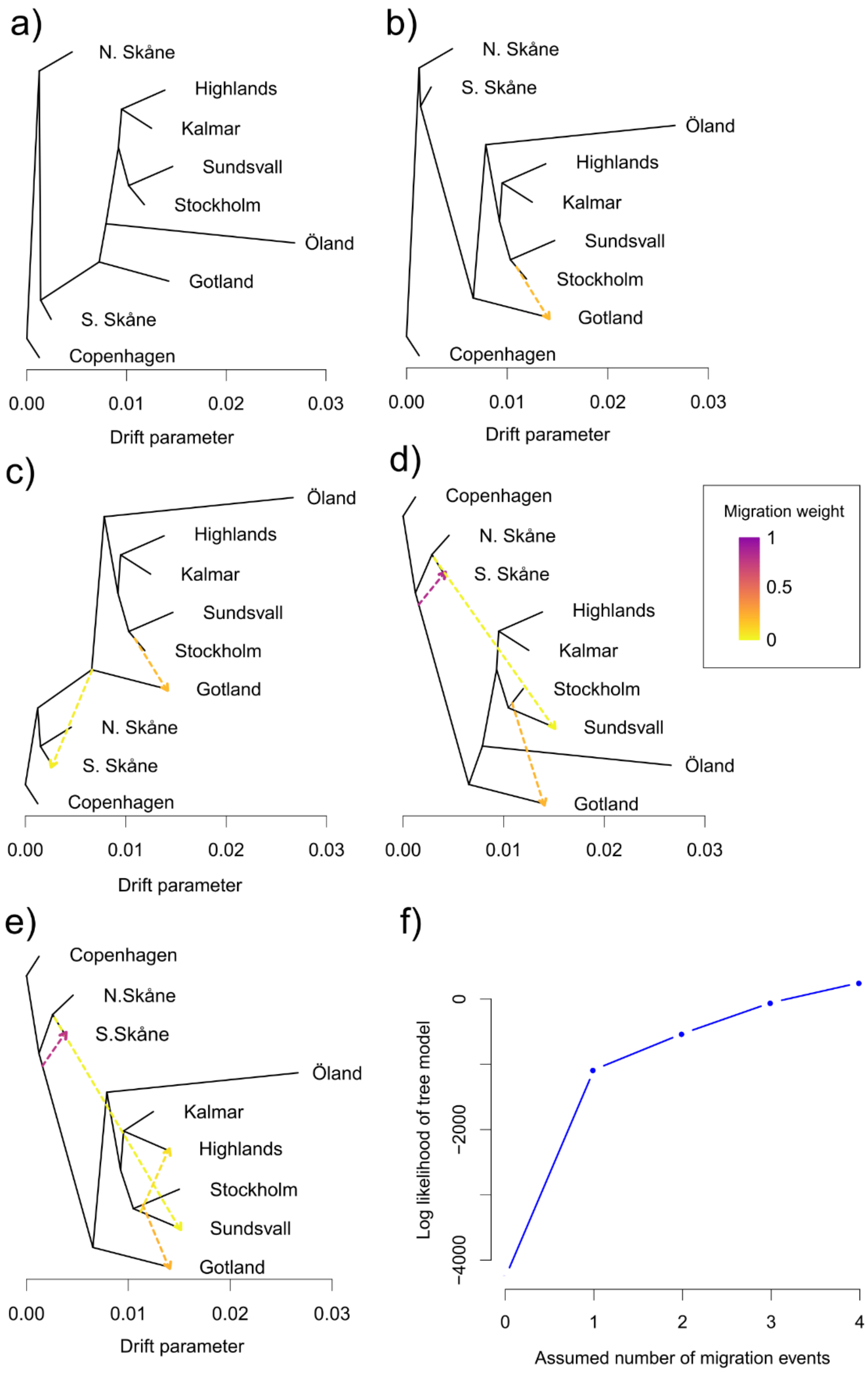


**Figure S2**. Phylogeographic relationships as inferred by *TreeMix* assuming no migration (a), or 1-4 migration events (b-e). Branch lengths represent the amount of genetic drift. Arrows represent inferred migration events; arrow color represents migration weight, i.e. the inferred proportion of the recipient population's ancestry that derives from the donor population. f) Log likelihood of tree model for each of the runs shown in a) through e). The *TreeMix* run illustrated in c) is the one reported in the main text.

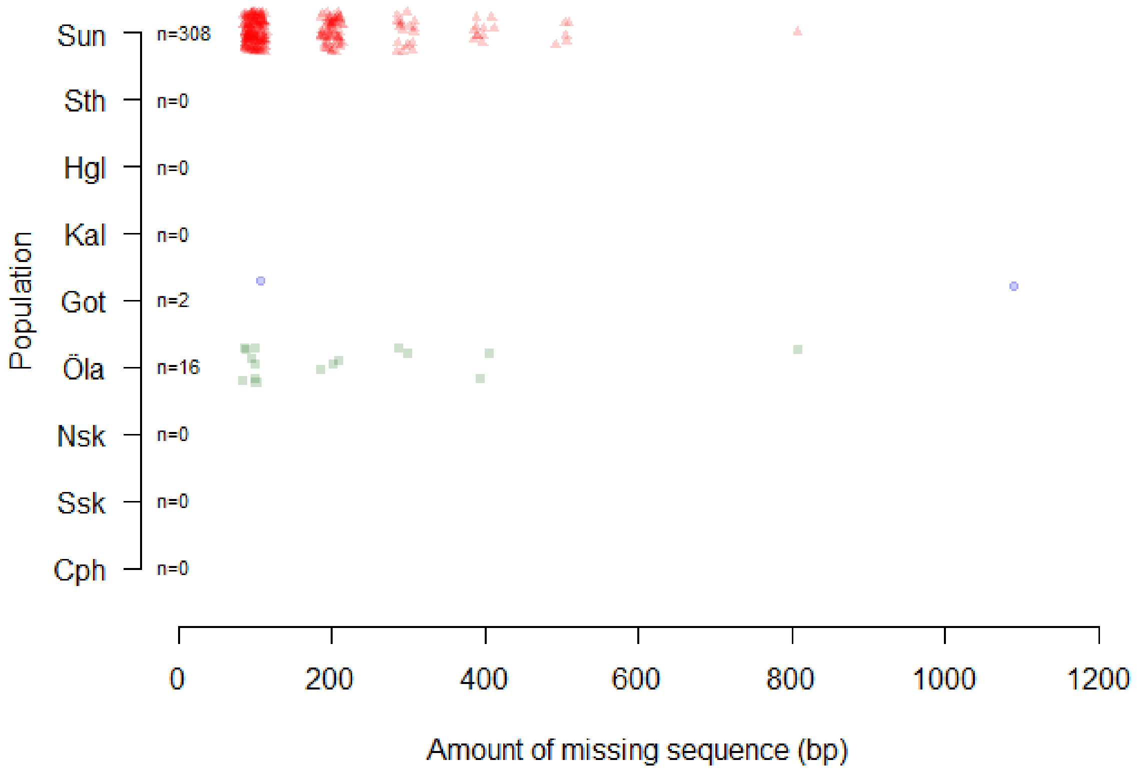


**Figure S3.** Counts and sizes of putative population-specific deletions across the *P. aegeria* genome. Each data point is a contig where at least one zero-read 100-bp window was found; x-axis value corresponds to the number of zero-read windows found in that contig. The Gotland-specific deletion in *timeless* was the largest deletion found.

**Table S1.** Details on population pools: pool size (number of haploid genomes=2N) and raw read depth statistics. During pileup subsampling, the 5$^{th}$ and 99$^{th}$ percentiles of raw read depth were used as the target and maximum read depth, respectively.

| | **Haploid pool size** | **5$^{th}$ perc. / target RD** | **99$^{th}$ perc. / max RD** | **Median RD** |
|---|---|---|---|---|
| Sundsvall | 44 | 14 (→ 20) | 51 | 32 |
| Stockholm | 60 | 23 | 67 | 46 |
| Highlands | 44 | 28 | 87 | 60 |
| Kalmar | 26 | 20 | 62 | 42 |
| Gotland | 56 | 36 | 102 | 72 |
| Öland | 58 | 46 | 129 | 95 |
| N. Skåne | 36 | 22 | 64 | 43 |
| S. Skåne | 88 | 48 | 134 | 98 |
| Copenhagen | 34 | 27 | 80 | 56 |
